# Host factor PLAC8 is required for pancreas infection by SARS-CoV-2

**DOI:** 10.1101/2023.08.18.553908

**Authors:** Lesly Ibargüen-González, Sandra Heller, Marta L. DeDiego, Darío López-García, Alba M Gómez-Valero, Thomas FE Barth, Patricia Gallego, Israel Fernández-Cadenas, Sayoa Alzate-Piñol, Catalina Crespí, Julieth A Mena-Guerrero, Eugenia Cisneros-Barroso, Alejandro P. Ugalde, Gabriel Bretones, Charlotte Steenblock, Alexander Kleger, Carles Barceló

## Abstract

**Background:** Although COVID-19 initially caused great concern about respiratory symptoms, mounting evidence shows that also the pancreas is productively infected by SARS-CoV-2. However, the severity of pancreatic SARS-CoV-2 infection and its pathophysiology are still under debate. Here we investigated the consequences of SARS-CoV-2 pancreatic infection and the role of the host factor Placenta-associated protein (PLAC8)

**Methods:** We analyzed plasma levels of pancreatic enzymes and inflammatory markers in a retrospective cohort study of 120 COVID-19 patients distributed in 3 severity-stratified groups. We studied the expression of SARS-CoV-2 and PLAC8 in the pancreas of deceased COVID-19 patients as well as in non-infected donors. We performed infection experiments in PLAC8 knock-out PDAC cell lines with full SARS-CoV-2 virus.

**Results:** We found that analysis of circulating pancreatic enzymes aided the stratification of patients according to COVID-19 severity and predict outcomes. Interestingly, we found an association between PLAC8 expression and SARS-CoV-2 infection in postmortem analysis of COVID-19 patients. Using full SARS-CoV-2 infectious virus inoculum from Wuhan-1 and BA.1 strains, we demonstrated that PLAC8 is necessary for productive infection of PDAC cell lines. Finally, we observed an overlap between PLAC8 and SARS-CoV-2 immunoreactivities of the pancreas of deceased patients.

**Conclusion:** Our data indicate the human pancreas as a SARS-CoV-2 target with plausible signs of injury and demonstrate that the host factor PLAC8 is required for SARS-CoV-2 pancreatic infection, thus defining new target opportunities for COVID-19-associated pancreatic pathogenesis.

**Plain language summary:** Previous studies have shown that the pancreas is infected by SARS-CoV-2. However, none of these studies have described measurable pancreatic damage associated to COVID-19 severity and the pathogenesis of pancreatic SARS-CoV-2 infection remains largely unknown. Novel host factors have been proposed for SARS-CoV-2 infection of mainly the airway epithelium, none of them studied in the pancreas.

Our study shows clinically relevant pancreatic damage associated with SARS-CoV-2 infiltration and assesses the predictive potential of circulating pancreatic enzymes to stratify patients according to COVID-19 severity and predict clinical outcomes in a cohort of 120 patients. Our data show that host factor Placenta-associated protein 8 (PLAC8) expression is linked to SARS-CoV-2 infection in postmortem analysis of COVID-19 patients and functionally demonstrated the full requirement of PLAC8 for SARS-CoV-2 pancreatic infection and viral replication.

Our data confirm the human pancreas as a SARS-CoV-2 target with signs of injury unveiling the measurement of pancreatic enzymes for prognosis value and demonstrating that host factor PLAC8 is required for SARS-CoV-2 pancreatic infection defining new stratification and target opportunities for COVID-19-associated pancreatic pathogenesis.

## Introduction

COVID-19, the respiratory disease caused by SARS-CoV-2, has been associated with a wide range of symptoms and complications, affecting not only the lung but also other organs in the body including the gastrointestinal system^1^ ^2^. Among the less studied, but potentially occurring with severe complications, is the infection of the pancreas, a gland responsible for producing hormones like insulin and digestive enzymes. While the exact mechanism of how SARS-CoV-2 affects the pancreas is not yet fully understood, several studies reported pancreatic inflammation, injury, and dysfunction in some COVID-19 patients^3–8^. Taken together, this raises concerns about the potential long-term impact on pancreatic function and the overall metabolic health of those affected.

Placenta-associated 8 (PLAC8) is a highly conserved protein that is expressed in various tissues like the placenta as well as the respiratory system or the gastrointestinal tract^9^. PLAC8 has been implicated in the regulation of multiple cellular processes like autophagy or cell motility^10^. Previous studies have suggested that PLAC8 also plays a prominent role in pancreatic neoplastic transformation^11–13^. Furthermore, PLAC8 has recently been identified as an essential host factor for SARS-CoV-2 infection in human lung cancer cell lines^14^ and for swine acute diarrhea syndrome coronavirus (SADS-CoV) in a swine primary intestinal epithelial culture^15^ supporting a potential role of PLAC8 as a pan-CoV infection factor.

In this work, we investigated the impact of SARS-CoV-2 infection on the pancreas as well as the potential relationship with PLAC8. Our data confirm previous studies reporting morphological changes associated with SARS-CoV-2 viral infiltration in the autopsy of the pancreas of patients who died from COVID-19. Notably, the quantification of circulating pancreatic lipase contributed to the stratification of patients according to COVID-19 severity. Interestingly, PLAC8 overexpression was associated with SARS-CoV-2 infection by postmortem analysis of COVID-19 patients. Functional studies employing pancreatic cancer cell lines revealed the requirement of PLAC8 for an efficacious infection of the pancreas. Finally, we observed an overlap in PLAC8 and SARS-CoV-2 immunoreactivity in the islets of deceased patients. In conclusion, our data confirm the human pancreas as a SARS-CoV-2 target and demonstrate that host factor PLAC8 is required for SARS-CoV-2 pancreatic infection defining new target opportunities for COVID-19-associated pancreatic pathogenesis.

## Methods

### Patients and samples. Ethics

#### Patient tissue for immunohistochemical analysis

Sections of human tissue from x patient (Patient ID: S27-20, Age: 89, Sex: female, comments: Died with COVID-19 in 2020)^6^ were provided by the pathology department of Ulm University. A board-certified pathologist (Thomas F.E. Barth) approved non-neoplastic tissue integrity. All experiments were executed according to the guidelines of the Ethics Committee of the Federal General Medical Council and approved by the Ethics Committee of the University of Ulm (vote for usage of archived human material 03/2014).

#### Patient tissue for immunofluorescence analysis

Autoptic material was retrieved from one patient who died of aortic dissection and from seven patients who died of COVID-19 in 2020-21. Pancreatic tissue was fixed in formalin and embedded in paraffin. The autopsies were performed at the Institute of Pathology at Universitätsklinikum Carl Gustav Carus in Dresden, Germany, or the Institute of Pathology at Universität Regensburg, Germany. Tissue samples were received from the pathology institutes with the respective approval of local ethics committees and collected in the frame of the German Registry for COVID-19 Autopsies (DeRegCOVID). Written informed consent to perform autopsies and to use the tissues for research purposes were obtained from patient relatives. Normal pancreatic tissues were obtained from Zyagen.

#### Samples for blood analysis

We performed a retrospective cohort study from a total of 80 subjects with COVID-19 that had a positive SARS-CoV-2 RT-PCR for whom blood samples were taken within 48 h of hospital admission between August 7th, 2020 and June 17th, 2021 obtained from the Biobank Unit located in the Hospital Universitari Son Espases (HUSE) or COVID19-negative healthy volunteers from the Blood and Tissue Bank of the Balearic Islands in the same period. This study was conducted in agreement with the Good Clinical Practice principles and the Declaration of Helsinki for ethical research. Ethical approval for this project (IB 4172-20 PI, 26 May 2020) was obtained from the ethics committee of the Balearic Islands and waived the requirement for informed consent, due to the emergency situation.

### Generation of PLAC8 knock-out cell lines

#### Plasmids

The plasmids pLenti-CRISPRv2-PLAC8-sg1 (KO1), -sg2 (KO2), used to generate PLAC8 KO cells, and the plasmid encoding a nontargeting control -NT sgRNA, were kindly donated by Prof Carlos López-Otín^14^.

#### Cell culture

The cell line SUIT-2 were provided by Dr. Marcos Malumbres (CNIO, Spain). African green monkey kidney epithelial Vero E6 cells were kindly provided by Prof. Luis Enjuanes (Centro Nacional de Biotecnología, CNB-CSIC, Spain). SUIT-2 was cultured in RPMI-1640 medium (Roswell Park Memorial Institute; Gibco), while HEK-293T and Vero E6 cells were cultured in DMEM medium (Dulbecco’s Modified Eagle Medium; Gibco). Both media were supplemented with 10% (v/v) (Gibco), 100 μg/ml of normocin (Invitrogen), 1 M HEPES (Gibco) and 1 % (v/v) of GlutaMax (Gibco). The cells were kept in an incubator at 37°C in a humid atmosphere with 5% CO_2_. Periodic passages were made using DPBS 1X (Dulbecco’s Phosphate Buffered Saline; Gibco) to wash the cells and TrypLETM Express 1X (Gibco) or EDTA-trypsin (Gibco) to recover them, since all the cell lines were attached and grew in monolayer. Tests were performed to detect contamination by *Mycoplasma sp*. by PCR every other week.

#### Lentivirus production for the generation of KO cells

To make a Knock-out (KO) cell, lentiviruses were produced with the different vectors of interest, following the recommendations of Invitrogen. 6×10^6^ HEK-293T cells were seeded in 10 cm plates (Sarstedt). The cells were transfected with 6 μg of psPAX2, 3.3 μg of 4 pMD2.G, and 7.7 μg of the corresponding pLenti plasmid (ratio 1.8:1:2.3 respectively), using Lipofectamine 3000 (Invitrogen). The following pLenti were used: pLenti-CMV-ZSGreen-P2AHygro as selection control, pLentiCRISPR v2 NT_0569 as a vacuum vector, and two pLentiRISPR v2 of two guides (sg1 and sg2) of PLAC8 for the realization of KO. After 6 h of incubation, the medium was replaced by Opti-MEM (Gibco) supplemented with GlutaMAX 1X, 1 mM sodium pyruvate (Gibco), and 5% FBS. At 24 h and 52 h post-transfection, viral supernatants were collected, filtered through a 0.45 μ m filter, and concentrated with PierceTM Protein Concentrator PES 50K MWC0 (ThermoFisher Scientific) at 14,000 g at room temperature up to 2 ml per plate. Finally, they were stored at −80°C until use.

#### Generation of PLAC8 KO cell lines

To generate a stable PLAC8 knock-out, 8×10^5^ cells per well were seeded in 24-well plates The next day, infections with lentiviruses containing pLenti-CRISPRv2-PLAC8-sg1 (KO1), -sg2 (KO2) and nontargeting control (NT) were performed using 200 μL of a 1:1 mixture of concentrated virus and media viral supernatants. At 48 h post-infection the selective medium was replaced: supplemented RPMI plus 2 μg/ml of puromycin. The selection was allowed for 10 days until all cells of a noninfected dish had died. At this point NT, KO1, and KO2 cell lines were pooled and knock-out was analyzed by Western Blot.

#### Western Blot analysis

To confirm the knock-out of PLAC8, cells were lysed in RIPA buffer containing protease and phosphatase inhibitors cocktail (Thermo Scientific, #78442). Protein concentration was determined using Pierce® BCA Protein Assay Kit (Thermo Scientific). Equal amounts of protein (20-30 μg) were resolved by 8 to 15% SDS PAGE MiniPROTEAN^®^ TGX gels (Bio-Rad), from samples previously prepared with Laemmli Buffer 4X™ (Bio-Rad) supplemented with 5% of β-mercaptoethanol (Sigma) and heated to 98°C for 5 min and transferred to Trans-Blot PVDF Transfer membranes (Bio-Rad, #1704157). Membranes were blocked for 1 h at room temperature with TBS-T (0.1% Tween 20) containing 10% milk or 5% BSA. Membranes were incubated at 4°C with primary antibodies diluted in TBS-T containing 10% milk:, anti-PLAC8 1:1,000 (HPA040465, Sigma), anti-ACE2 1:1,000 (SN0754, Invitrogen), anti-Actin Monoclonal 1:1,000 (ACTN05, Invitrogen), anti-beta-Actin Polyclonal 1:2,000 (20536-1-AP, Proteintech), anti-Alpha-Tubulin Polyclonal 1:1,000 (11224-1-AP, Proteintech), and anti-GFP (SAB4301138, Sigma-Aldrich). After washing with TBS-T, the membranes were incubated for 2 h at room temperature with secondary antibodies in TBS-T containing 10% milk: anti-Mouse IgG (HAF018, R&D Systems) and anti-Rabbit IgG (A9196, Sigma-Aldrich) at 1:2,000 and 1:10,000 dilutions, respectively, both conjugated with HRP. Finally, the protein bands were visualized using Clarity^TM^ Western Blot ECL (Bio-Rad) and recorded with ImageQuant LAS4000 (Fujifilm).

### SARS-CoV-2 infection assays

#### Viruses and virus titrations

Virus stocks of SARS-CoV-2, Wuhan-1 and BA.1 strains, were grown in Vero E6 cells, respectively, under BSL3 conditions. Both strains (kindly provided by L. Enjuanes, CNB-CSIC) were isolated in Vero E6 cells starting from nasopharyngeal swabs from patients with COVID-19 from Hospital 12 de Octubre, Madrid, Spain^40^ and unpublished data.

SARS-CoV-2 virus titrations were performed in Vero E6 cells grown in 24-well plates and infected with ten-fold serial dilutions of the virus, as previously described^41^. After 1 h absorption, cells were overlaid with low electroendosmosis agarose (Pronadisa) and incubated for 3 days at 37 °C. For all virus titrations, cells were fixed with 10% formaldehyde in phosphate buffer saline (PBS) and permeabilized with 20% methanol. Viral plaques were visualized and counted using crystal violet staining.

#### Immunofluorescence microscopy and flow cytometry

Confluent monolayers of human SUIT-2 cells grown on sterile glass coverslips (24-well plate format) were mock-infected or infected (MOI 1) with SARS-CoV-2 (Wuhan-1 and BA.1 strains). At 24 hours post-infection (hpi), the cells were fixed and permeabilized with 10% formaldehyde and 0.1% Triton X100 for 20 min at room temperature. Then, cells were blocked with 2.5% BSA in PBS and SARS-CoV-2 nucleocapsid (N) protein was detected with a rabbit anti-N antibody at 1:1000 dilution (Genetex GTX135357). Coverslips were washed with PBS for 4 times, and incubated with secondary anti-rabbit Ab conjugated to Alexa Fluor 488 (Invitrogen), for 45 min at room temperature. Nuclei were stained using DAPI (ThermoFisher Scientific). Coverslips were analyzed on a Leica DMi8 S widefield epifluorescence microscope. Images were acquired with the same instrument settings and analyzed using the Fiji software. To calculate the percentage of infection, more than 600 cells per condition were counted.

Alternatively, confluent monolayers of human SUIT-2 cells grown on sterile glass coverslips (6-well plate format) were mock-infected or infected (MOI 1) with SARS-CoV-2 (Wuhan-1 and BA.1 strains). At 24 hours post-infection (hpi), the cells were detached from the wells and the pelleted cells were fixed and permeabilized with 10% formaldehyde and 0.1% saponin for 20 min at room temperature. Then, cells were blocked with 10% FBS in PBS and SARS-CoV-2 nucleocapsid (N) protein was detected with a rabbit anti-N antibody (Genetex GTX135357). Coverslips were washed with PBS and 0.1% saponin for 4 times, and incubated with secondary anti-rabbit Ab conjugated to Alexa Fluor 488 (Invitrogen), during 45 min at room temperature. The cells were analyzed in a FACSAria Fusion (Becton Dickinson), and the percentage of infected cells, was determined using the FlowJo software (TreeStar).

### Blood analysis

#### Patients

We performed a retrospective cohort study from a total of 80 subjects with COVID-19 that had a positive SARS-CoV-2 RT-PCR for whom blood samples were taken within 48 h of hospital admission between August 7th, 2020 and June 17th, 2021 obtained from the Biobank Unit located in the Hospital Universitari Son Espases (HUSE) or COVID19-negative healthy volunteers from the Blood and Tissue Bank of the Balearic Islands in the same period. This study was conducted in agreement with the Good Clinical Practice principles and the Declaration of Helsinki for ethical research. Ethical approval for this project (IB 4172-20 PI, 26 May 2020) was obtained from the ethics committee of the Balearic Islands and waived the requirement for informed consent, due to the emergency situation.

Hence, three patient subgroups of 40 sex and age-matched patients were defined using the WHO ordinal scale of clinical improvement (OSCI) as described^19^: 40 control cases (WHO OSCI=0), 40 patients with severe COVID-19 (WHO OSCI=3–4) admitted to the Ward, and 40 patients with critical COVID-19 (WHO OSCI=5–8) admitted to the ICU. Our control group comprised 40 healthy uninfected subjects (Healthy), matched by age and sex.

#### Plasma samples measurements

Plasma samples were collected in two 10-ml EDTA and a 4.5-ml tube with sodium citrate and centrifuged at 2,000×g for 10 min as previously described (Durat et al, 2014) and fractionated by automation (Janus robotic liquid handlers; PerkinElmer). Typically, 6-8 h elapsed between sample collection and freezing.

Clinical data were collected during the patient standard of care and final discharge diagnosis. Measurements of C-Reactive protein (CRP), and D-Dimer, Lipase and Protease plasma concentrations were determined at Hospital Son Espases Facilities through the hospital automation system by Alinity I (AbbottDiagnostics).

### Immunofluorescence analysis

#### Immunofluorescence

Paraffin sections were deparaffinized in Neo-Clear (Merck) and rehydrated through a descending graded ethanol series. Antigen retrieval was performed in citrate retrieval buffer pH 6.0, using a Decloaking Chamber NXGEN (Menarini Diagnostics) at 110°C for 3 mins. Sections were blocked in blocking buffer (PBS containing 1% BSA (w/v), 0.1% (v/v) Triton X-100, 5% (v/v) goat serum) for 1 h at room temperature, followed by incubation with primary antibody diluted in PBS containing 1% BSA (w/v), 5% (v/v) goat serum at 4°C overnight. Slides were washed in PBS and incubated with appropriate fluorophore-conjugated secondary antibodies in PBS for 2 h at room temperature. Slides were washed in PBS and nuclei were stained with 4’-6-diamidino-2-phenylindole (DAPI; Thermo Fisher Scientific) and mounted with an aqueous mounting medium (Aqua-Poly/Mount; Polysciences).

#### Confocal laser scanning microscopy and fluorescence microscopy

Confocal imaging was performed with a Zeiss LSM 880 inverted confocal laser scanning microscope and ZEN 2010 software (Zeiss). Image processing and analysis were carried out using ImageJ version 1.53q software. The mean fluorescence was measured using ImageJ.

### Immunohistochemical analysis

#### PLAC8 and Spike staining

Tissue sections were deparaffinized, rehydrated and subjected to heat-mediated antigen retrieval in citrate buffer, pH 6. After blocking of endogenous peroxidases (3% H_2_O_2_) and unspecific antibody binding (2% BSA+5%serum) PLAC8 antibody (HPA040465, Sigma) or SARS-CoV-2 Spike Glycoprotein S1 antibody (GTX632604, Biozol) where incubated overnight at 4°C. Detection was performed by VECTASTAIN^®^ Elite^®^ ABC-HRP and NovaRED^®^ Substrate (Vector laboratories).

#### SARS-CoV-2-N antibody validation

For the immunofluorescence stainings of SARS-CoV-2 in pancreatic tissue, a mouse monoclonal antibody against the SARS-CoV-2 nucleocapsid protein (SARS-CoV-2-N, Sino Biological, 40143-MM05) was used^5^. This antibody was validated using negative control pancreatic tissue and lung tissue from a COVID-19 patient that was previously shown to be positive for both the SARS-CoV-2 spike 1 and the nucleocapsid 2 protein. In lung tissue from a COVID-19 patient, many infected cells were detected with some background fluorescence (Fig. S1A). In pancreatic negative control tissue (non-COVID-19), there was no signal, whereas the antibody exhibited a clearly positive signal in tissues from COVID-19 patients (Fig. S1B). Therefore, this antibody was used for all immunofluorescence stainings.

#### PLAC8 quantification

Bright-field images were acquired using a Leica DM5500B microscope (Leica) with a Leica DMC5400 camera and Leica Application Suite software. Measurement of PLAC8 stained area was performed using QuPath Software (Bankhead, P. et al. QuPath: Open source software for digital pathology image analysis^39^

Briefly, NovaRED and hematoxylin stainings have been separated using the color deconvolution method followed by thresholding stained areas and counting of pixels in positive classifications. Additionally, areas without tissue have been excluded by setting a threshold or by manual annotation when necessary. Regions of interest containing Islets of Langerhans and Acini have been annotated manually. Accordingly, PLAC8 positively-stained area per total tissue area or annotated area has been calculated.

### Mendelian randomization

Mendelian randomization (MR) is a statistical method that leverages genetic variations, specifically single nucleotide polymorphisms (SNPs), to assess causality^27^. The main MR method was inverse-variance weighted. Also, complementary MR approaches were applied: MR-Egger, weighted median, penalized weighted median, and weighted mode approaches. Statistical analysis was conducted in R using TwoSampleMR (version 0.5.6. Stephen Burgess, Chicago, IL, USA). The methodology followed in this study was in accordance with the latest guidelines for Mendelian randomization studies.

To evaluate these causal effects, we use free access summary statistics data from COVID19-hg Genome Wide Association Studies (GWAS) meta-analyses of three different phenotypes available at https://www.covid19hg.org: very severe respiratory confirmed COVID-19 vs. population, hospitalized COVID-19 vs. population cohorts and COVID-19 vs. population cohorts.

The data included studies of populations of various genetic ancestries, including European, mixed-race Americans, Africans, Middle Eastern, South Asian and East Asian individuals. Severe COVID-19 cases were defined as those individuals who required hospital respiratory care or died from the disease, using 9,376 cases and 1,776,645 control individuals in this cohort. Moderate or severe COVID-19 cases were defined as those participants who were hospitalized due to symptoms associated with infection in this cohort using 25,027 cases and 2,836,272 control individuals. And finally, a cohort of all cases with SARS-CoV-2 infection reported regardless of symptoms was used and included 125,584 cases and 2,575,347 control individuals. Controls for all three analyses were selected as genetically matched samples by ancestry with no known SARS-CoV-2 infection^14,40^. On the other hand, summary statistics for each enzyme used for MR have been obtained from DECODE GENETICS available at www.decode.com/summarydata/ from the Icelandic Cancer Project using a cohort of almost 40,000 patients and a control population, randomly selected from the National Registry^42^.

Significantly associated (p-value < 5×10^−8^) single nucleotide polymorphisms (SNPs) for each enzyme were used as instruments for MR analysis and then performed a clumping of the variants with a threshold of r2 < 0.001 using the 1000 G European reference panel. For the significant and robust results, we performed scatterplots and leave-one-out tests that are shown in forest plots. A significant causal association was considered to exist when the p-value was lower than 0.05, complementary methods showed the same direction of the association and no pleiotropy or heterogeneity was detected.

### Statistical analysis

Statistical analysis was performed using GraphPad Prism version 9.0.0 (GraphPad Software, San Diego, CA, USA). Statistical significance was determined using an unpaired two-tailed Student’s *t*-test for comparison of two means. The significance was defined as*: ns (not significant) P > 0.05; * P < 0.05; ** P < 0.01; *** P < 0.001; **** P < 0.0001*.

## Results

### Pancreatic damage is associated with COVID-19 severity

Given the rather open debate about the magnitude of COVID-19 impact on pancreatic tissue^5,6,16,17^, we sought to investigate the damages inflicted by the SARS-CoV-2 virus in the infected patients featuring different degrees of disease severity.

First, we studied postmortem material from patients who died from COVID-19. We stained sections for viral SARS-CoV-2 Spike protein, previously validated for immunohistochemistry (**Fig S1**) in a SARS-CoV-2 infected placenta and lung as well as infected 293T cells. In accordance with previous studies^6,18^, pancreatic histopathology revealed the presence of SARS-CoV-2 Spike protein in islets and epithelium of the acini. This indicates a persistent productive infection during severe COVID-19 (**Fig 1A**).

**Fig 1.-.**
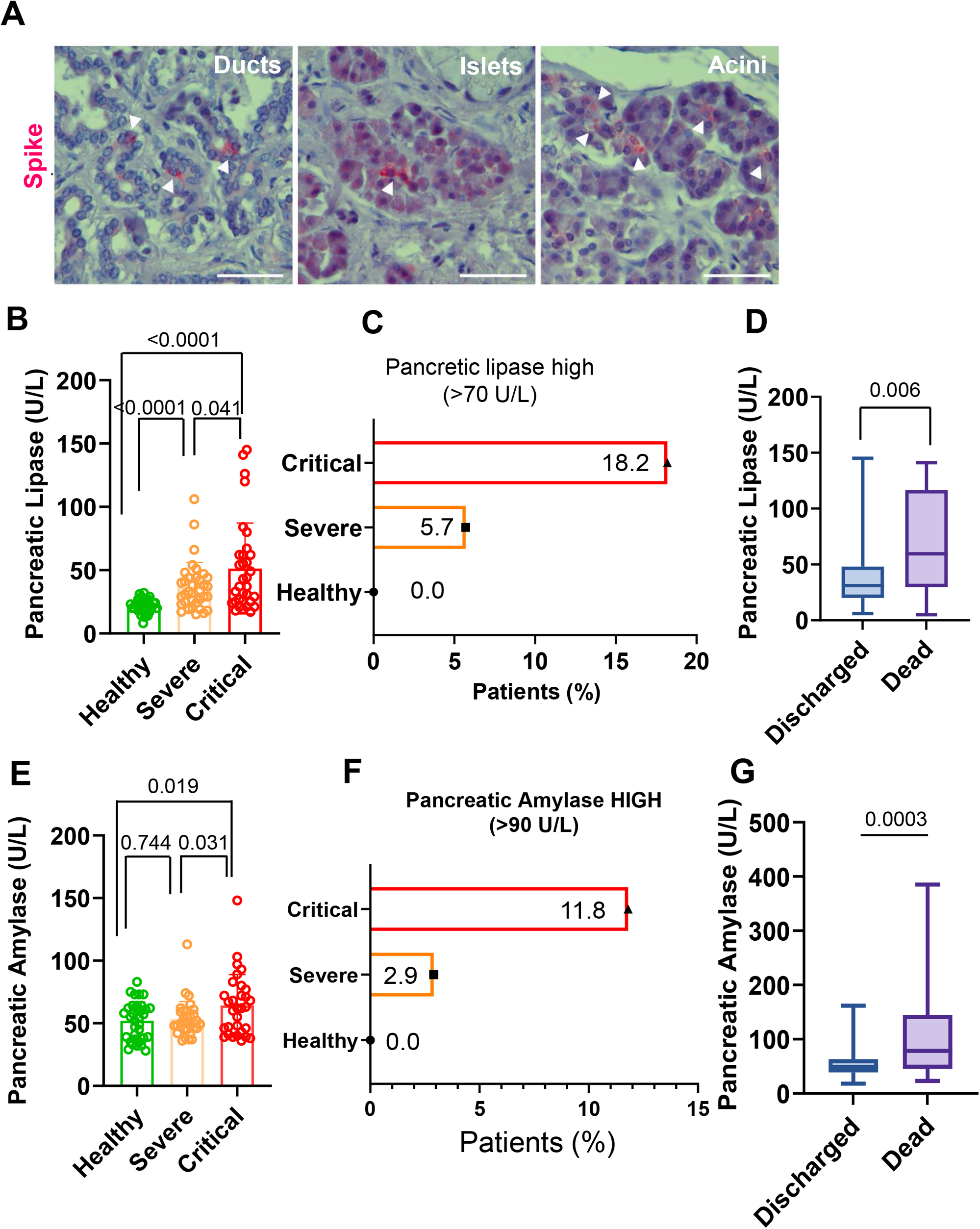
SARS-CoV-2 pancreatic infection is associated with damage. **A)** Viral infiltration occurs in both endocrine (Islets) and exocrine pancreas (Ducts and Acini). Sections are stained with anti-SARS-CoV-2 Spike Glycoprotein S1 (Spike,red) and counterstained with Hematoxylin. Bar, 100 µm. **B)** Plasma levels of pancreatic injury biomarker pancreatic lipase stratify COVID-19 patients according to severity; **C)** Patients (%) with pancreatic lipase above normal blood levels (>70 U/L); **D)** Plasmatic levels of pancreatic lipase in hospitalized patients that ultimately were discharged or who died; **E)** Plasma levels of pancreatic injury biomarker pancreatic amylase stratify COVID19 patients according to severity**; F)** Patients (%) with circulating pancreatic amylase above normal blood levels (>90 U/L); **G)** Plasmatic levels of pancreatic lipase in hospitalized patients that ultimately were discharged or who died.

As previously described^18^, the morphology of some of the infected cells did not resemble ductal, acinar, or endocrine architecture, indicating a certain degree of tissue hyperplasticity occurring after infection (e.g. shrinkage of the islets, degeneration, and polyploidy of the β-cells were observed). In some patients, mild lymphocytic infiltration of both the endocrine and exocrine pancreas was demonstrated (**Fig S1**).

To complement these findings, we analyzed signs of pancreatic injury by measuring plasma levels of pancreatic injury-related enzymes. To this end, we studied plasma in a retrospective cohort study of 120 patients classified into three severity-stratified groups according to the WHO ordinal scale of clinical improvement (OSCI)^19^: 40 control cases (WHO OSCI=0), 40 patients with severe COVID-19 (WHO OSCI=3–4), and 40 patients with critical COVID-19 (WHO OSCI=5–8).

Demographic and clinical characteristics of the three groups were stratified according to their severity assignments by the clinical care team at Hospital Son Espases (HUSE), Palma, Spain (**Table 1**). The study cohort comprised 47% females and featured an average age of 61.7 years. The most frequent co-morbidities were dyslipidemia (43.3%), obesity (30%), and hypertension (26.7%). Severe and critical severity groups were very similar for most characteristics and the control group exhibited significant differences with respect to Total (p<0.05) in Age (56 vs 61.7 y, p<0.0001), Hypertension (10% vs 26.7%, p=0.012), Type-2 Diabetes Mellitus (T2DM) (2.5% vs 19.3%, p=0.0039) and Chronic Obstructive Pulmonary Disease (COPD) (0% vs 9.2%, p=0.0031) (**Table 1**).

**Table 1.** Baseline characteristics of the sample population.

|  | **Total**<br>**(n=120)** | **Control**<br>**group**<br>**(n=40)** | **Severe**<br>**COVID19**<br>**(n=40)** | **Critical**<br>**COVID19**<br>**(n=40)** | **p-value** |
| --- | --- | --- | --- | --- | --- |
| *Age (years)* | 61.7 | 56 | 66.3 | 62.8 | <0.0001* |
| *Sex* |  |  |  |  | 0.1753 |
| *Female* | 56 (47%) | 22 (55%) | 20 (50%) | 14 (35%) |  |
| *Male* | 64 (53%) | 18 (45%) | 20 (50%) | 26 (65%) |  |
| **Comorbidities** |  |  |  |  |  |
| *Chronic pancreatitis* | 1 (0.83%) | 0 (0%) | 0 (0%) | 1 (2.5%) | 0.1308 |
| *Hypertension* | 32 (26.7%) | 4 (10%) | 13 (32.5%) | 15 (37.5%) | 0.0124 |
| *Type 2 DM* | 23 (19.3%) | 1 (2.5%) | 10 (25%) | 12 (30%) | 0.0039 |
| *Obesity* | 36 (30%) | 7 (17.5%) | 17 (42.5%) | 12 (30%) | 0.0510 |
| *COPD* | 11 (9.2%) | 0 (0%) | 4 (10%) | 7 (17.5%) | 0.0031 |
| *Dyslipidemia* | 52 (43.3%) | 9 (22.5%) | 23 (57.5%) | 20 (50%) | 0.0040 |
| *Smokers* | 18 (15%) | 3 (7.5%) | 5 (12.5%) | 10 (25%) | 0.0782 |

Data are presented as the mean (age), absolute numbers and percentage (%). \* One-way ANOVA was used to calculate the *P*-value. The rest are calculated with the Chi-square test

Elevated blood levels of pancreatic enzymes such as lipase and amylase are typically documented during pancreatic injury^20–22^. Given its rather long-lasting elevated levels, pancreatic lipase plasma levels not only render as the favored circulating diagnostic biomarker for pancreatic injury in terms of sensitivity and diagnostic window but is also a useful diagnostic biomarker in early and late stages of pancreatitis^23–25^

In our experimental setting, pancreatic lipase plasma levels aided to significantly stratifying patients according to severity in three groups (healthy vs severe and healthy vs critical, p<0.0001; severe vs critical, p=0.041) (**Fig 1B**). Significant discrimination between the critical and severe groups was also achieved upon measuring pancreatic amylase, a less specific biomarker for pancreatic damage (**Fig 1E**). Moreover, plasma levels of both pancreatic enzymes, indicative of pancreatic damage, exhibited a significant correlation between themselves (r=0.46, p<0.0001) (**Fig S2A**).

The normal blood level of amylase and lipase are 0–90 U/L and 0–70 U/L, respectively^22,26^. Interestingly, the number of patients classified according to the levels of these enzymes in moderate or elevated, mirrored the magnitude of COVID-19 disease. Patients with higher pancreatic lipase blood levels than baseline (hyperlipasemia) showed a 3-fold increase (**Fig 1C**) while patients with higher amylase levels exhibited a 4-fold increase (**Fig 1F**) when comparing critical vs severe COVID-19 patients. None of the healthy donors exhibited an increase of these enzymes (**Fig 1C, F**)

Notably, the differentiation of severity groups related to circulating pancreatic enzymes resembled the distribution of the inflammatory markers C-reactive protein (CRP) and D-Dimer that are hallmarks of severe COVID-19 (**Fig S2B, D).** Moreover, the Pearson correlation demonstrated that levels of plasma pancreatic lipase significantly correlated with these two typically used clinical variables that discriminate severity-stratified groups, CRP (r=0.27 p=0.018) and D-Dimer (r=0.47 p=0.004) (**Fig S2C, E**)

We then tested the potential ability of pancreatic lipase and amylase plasma levels in discriminating between patients who ultimately were discharged or who died in hospitalized centers. We found that both, plasmatic pancreatic lipase and amylase represent parameters that can reliably be used to significantly discriminate between these two groups (Pancreatic lipase, p=0.006; Amylase, p=0.0003) (**Fig 1D, G**) further reinforcing the role of pancreatic damage in COVID-19 pathogenesis.

Finally, we used a two-sample Mendelian Randomization (MR) approach^27^ to evaluate the causal association between COVID-19 severity and plasma levels of the enzymes pancreatic amylase and pancreatic lipase.

Along the same lines as the other results for pancreatic amylase, those patients with COVID-19 severe or critical symptoms had higher levels of biomarkers of pancreatic damage. We analyzed 8,779 cases with severe or critical COVID-19 symptoms and 1,001,875 controls^28^: odds ratio (OR), 0.95 ([95% CI, 0.91–1.01], standard error (SE)=0.07, beta=0.1 and p value=0.03). Although the leave-one-out analysis showed that causality was not driven by a single nucleotide variant, there was no pleiotropy or heterogeneity, and complementary methods showed the same direction of the association **(Fig S3A, B).** In contrast, no significant associations were observed with the inpatient and COVID-19 vs. population groups and no significant associations were observed with pancreatic lipase enzyme.

### PLAC8 is overexpressed in the SARS-CoV-2-infected pancreas

A potential link between SARS-CoV-2 infection and the upregulation of PLAC8 expression in lung epithelial cells has been reported^29^. Since PLAC8 has been described to be prominently expressed in the perturbed pancreas^10–12^, we analyzed the association between SARS-CoV-2 infection and PLAC8 distribution in pancreatic tissues from deceased individuals with COVID-19.

We studied the distribution of PLAC8 in the pancreas by immunohistochemical analysis of postmortem material from a patient described in Müller et al^6^ and found that PLAC8 was mainly expressed in islets and to a lesser extent in acini (**Fig 2A,B; Fig S4**). This is consistent with a putative role of PLAC8 during viral infection given the detection of significantly higher levels of this protein in SARS-CoV-2-infected patients (n=3) than in non-infected individuals (n=3) in the pancreatic tissue (p<0.0001), acini (p=0.002) and islets (p=0.0002) (**Fig 2A,B**). We substantiated our quantifications upon using an independent series of SARS-CoV-2 infected patients (n=2) vs non-infected individuals (n=3) and quantified the expression levels in immunofluorescence stainings. We obtained similar results for the pancreatic tissue (p=0.021) and islets (p=0.0017) while non-significant differences were documented in acini (p=0.167) (**Fig S5**).

**Fig 2.-.**
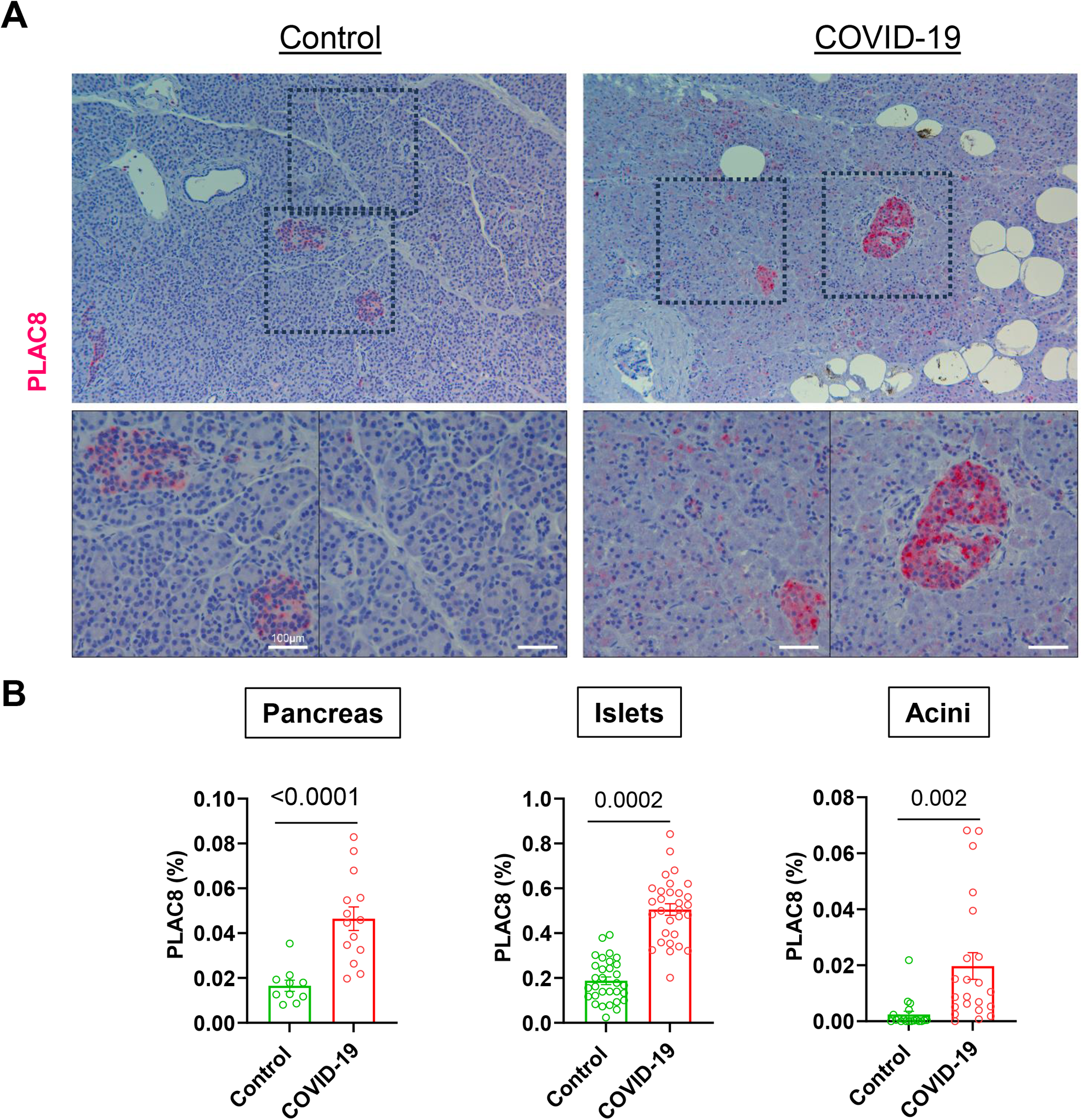
Host factor PLAC8 is overexpressed in the pancreas of COVID-19 deceased patients. **A)** PLAC8 expression pattern in pancreatic sections of non-infected (Control) and infected (COVID-19) deceased patients. Sections are stained for PLAC8 protein (red) and counterstained with Hematoxylin (blue). Rectangles mark areas of higher magnification in the bottom row. Bar, 100 µm; **B)** Pancreatic Islets and Acini were manually identified in necropsies and blinded quantified using Fiji. Data are presented as mean ± s.e.m. of PLAC8 positive area relative to total area for uninfected (Control), and infected (COVID-19). Statistical significance was calculated by ordinary two-tailed unmatched t-test

These results reiterate that PLAC8 could be implicated in pancreatic COVID-19 pathogenesis.

### SARS-CoV-2 productive infection is PLAC8-dependent in a pancreatic cancer cell line

To validate the effect of PLAC8 loss-of-function using full SARS-CoV-2 infectious viruses from Wuhan-1 and BA.1 strains, SUIT-2 PLAC8 NT and KO cell lines were infected with a SARS-CoV-2 inoculum. Viral infection efficacy was assessed by immunofluorescence of the nucleocapsid (N) protein at 24 h postinfection (p.i.). Consistent with our previous results, PLAC8 loss-of-function caused a significant reduction in the number of SARS-CoV-2 infected cells at 24 h p.i. in KO cell for both viral strains (Wuhan-1, p=0.0002; BA.1, p=0.0001).(**Fig 3A**).

**Fig 3:**
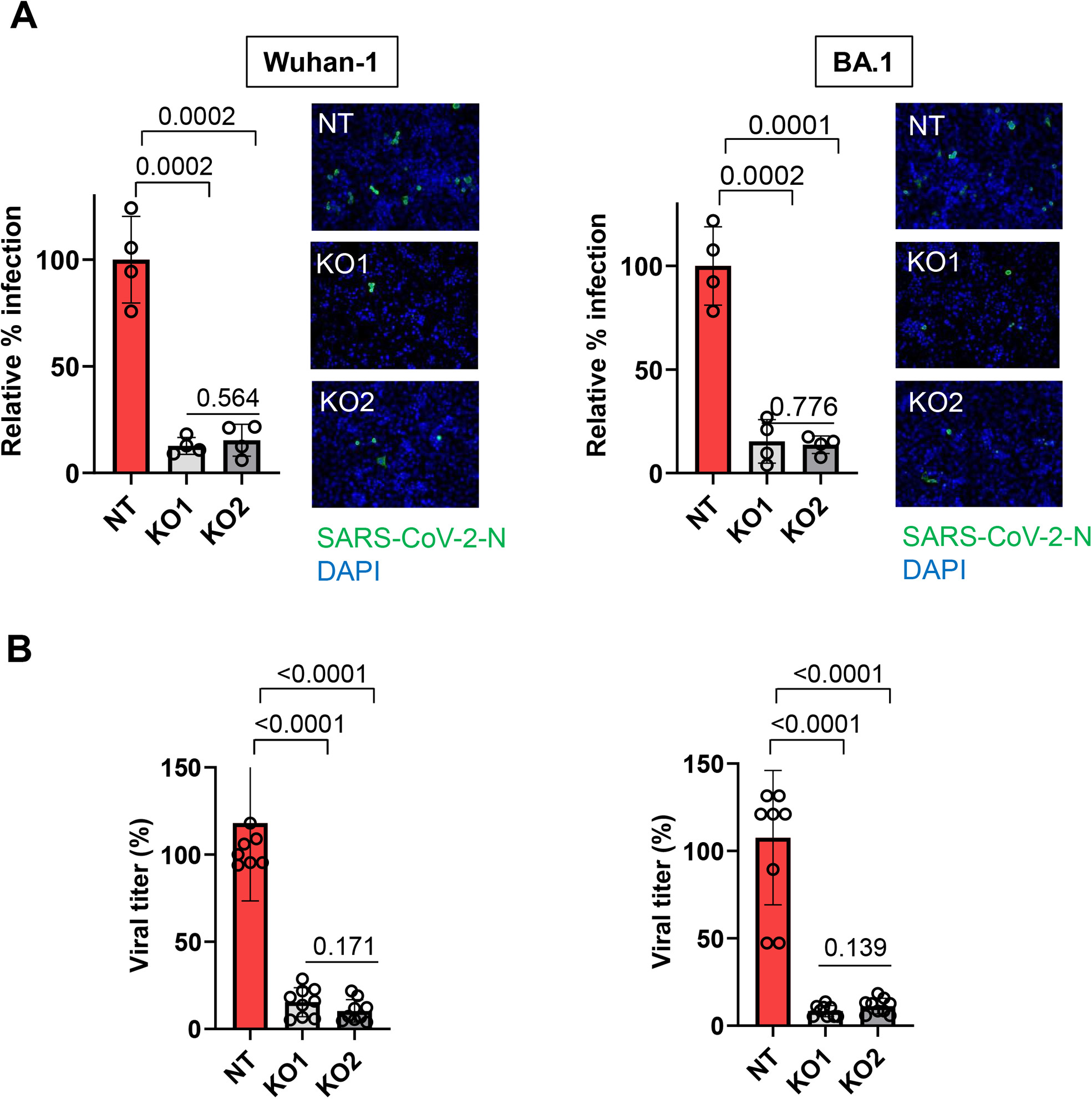
SARS-CoV-2 productively infects pancreatic cell lines in a PLAC8-dependent manner. **A)** Susceptibility to infection with two strains of full SARS-CoV-2 viruses in SUIT-2 cell line with loss-of-function of PLAC8. immunofluorescence quantification of the relative number of infected cells at 24 h post-infection in the PLAC8 loss-of-function cell lines relative to CRISPR non-targeting transduced (NT) control cells infected with SARS-CoV-2. Right: representative immunofluorescence images of SARS-CoV-2 nucleocapsid protein (green) and cell nuclei (blue) in the indicated cell models. **B)** Viral titer count of SARS-CoV-2 in SUIT-2 supernatant at 24 h postinfection. Bars represent the average and SEM of the percentage of infected cells or virus titers relative to CRISPR non-targeting transduced (NT) cells in three biological replicates (three different CRISPR KO cell lines infected independently). The significance above each bar represents the test p-value between each condition and CRISPR non-targeting transduced (NT) cells.

The viral replication capacity in SUIT-2 PLAC8 NT and KO cell lines was evaluated by analyzing infectious viral titers in the respective cell supernatants. Our results show that an efficient viral replication in the SUIT-2 cell line was significantly impaired by loss-of-function of PLAC8 for both viral strains (p<0.0001) (**Fig 3B**).

### Host factor PLAC8 co-localizes with SARS-CoV-2 during severe COVID-19

To further confirm PLAC8 requirement for SARS-CoV-2 infection event in pancreatic tissue, expression patterns of PLAC8 and SARS-CoV-2 were analyzed on autopsies from 2 individuals deceased from COVID-19. An archival control (Ctrl #1, non-infected pancreatic tissue collected in 2020) was used to ensure the specificity of SARS-CoV-2 nucleocapsid (N) protein (SARS-CoV-2N) staining. As expected, in this control, the SARS-CoV-2N signal was weak when compared to the SARS-CoV-2N staining in the COVID-19 patients (**Fig 4A,B**). Sections were subjected to the evaluation of not only PLAC8 but also insulin immunoreactivity to define the islet compartment. PLAC8 staining showed a stronger signal in islets as compared to acini in agreement with our previous observations (**Fig 2A**).

**Fig 4.-.**
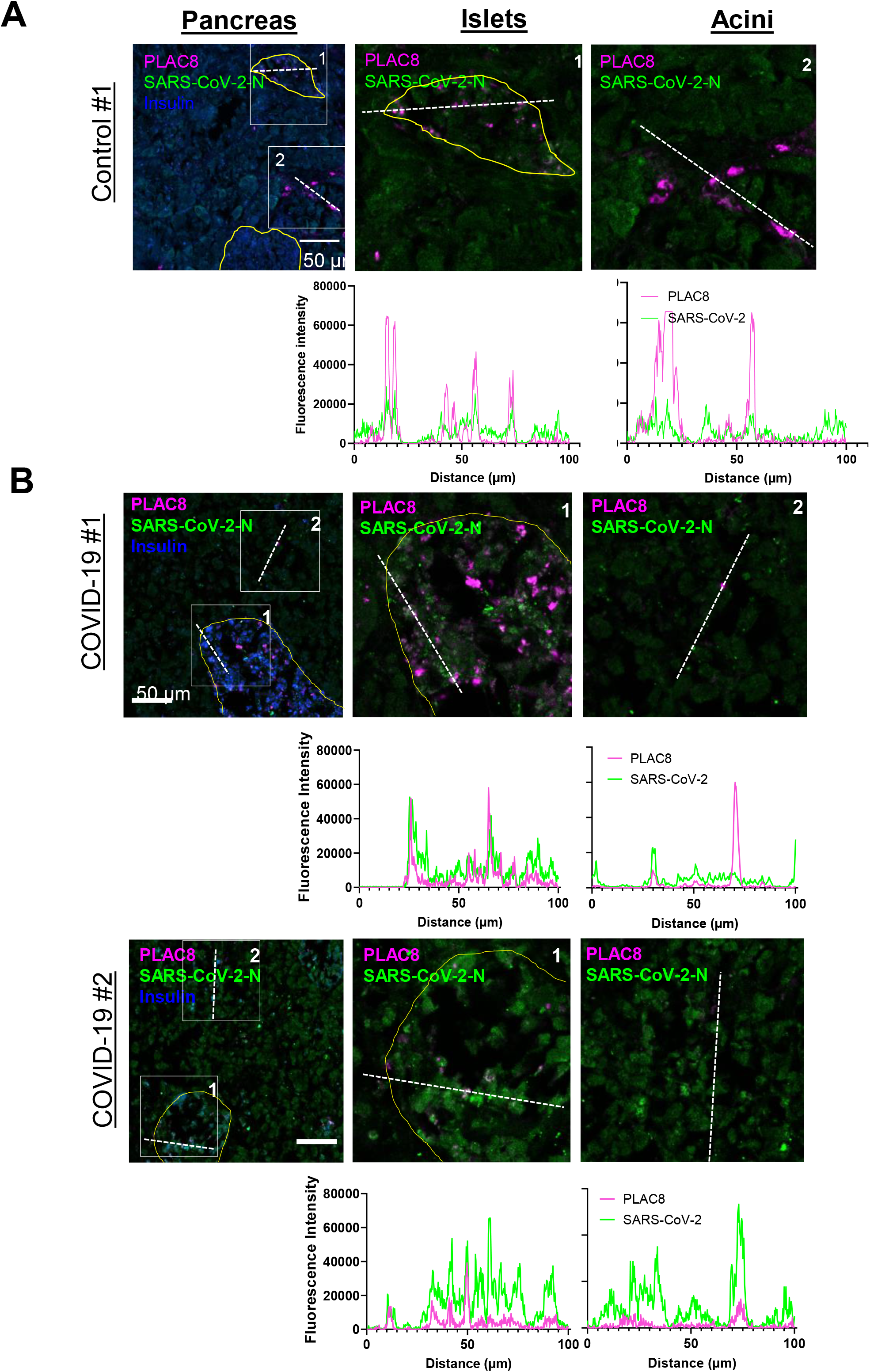
Host factor PLAC8 expression correlates with SARS-CoV-2 infection in pancreas of COVID-19 deceased patients. Morphometric analysis of a PLAC8/SARS-CoV-2-N staining of pancreatic tissue from non-infected controls **(A)** and COVID-19 patients **(B).** The fluorescence intensities in the indicated fluorescence channels along the white line were measured. Scale bar, 100 µm.

Morphometric analysis showed that PLAC8 co-localized with SARS-CoV-2N. A robust co-localization was documented especially in the islets, and to a lesser extent in the acinar compartment (**Fig 4B**).

Altogether, these experiments point to the requirement of host factor PLAC8 for efficient and productive infection.

## Discussion

Although COVID-19 initially caused great concern about respiratory symptoms, most patients also manifested gastrointestinal symptoms^1,2^ and mounting evidence shows that both the endocrine and exocrine pancreas are productively infected by SARS-CoV-2^5,6,8^. However, the prevalence and severity of pancreatic COVID-19 infection as well as its pathophysiology, are still under debate.

In agreement with previous observations^6,32^, IHC analysis of COVID-19 postmortem specimens revealed a substantial SARS-CoV-2 viral infiltration in both, endocrine and exocrine compartments of pancreatic tissue. Importantly, we could also observe morphological changes indicative of cell damage as previously reported for SARS-CoV-2^6,18^ and SARS-CoV^33^. In the previous pneumonia pandemic caused by SARS-CoV infection (2003), the virus was detected not only in the tissues of the lung, liver, kidney, and intestine but also of the pancreas, indicating the pancreas as a potential coronaviral target^34^. Moreover, SARS-CoV infection caused damage of the islets and subsequent acute diabetes^33^ suggesting that pancreas is a pan-CoV target with high clinical relevance for potential future CoV pandemics.

Case reports and retrospective cohort studies have revealed an association between pancreatitis-associated hyperlipasemia/hyperamylasemia and COVID-19^21,34^. As SARS-CoV-2 receptors are expressed in the pancreas and endothelial damage can occur, this association is indeed plausible^20^. To interrogate the pancreatic damage inflicted by the SARS-CoV-2 infection, we studied plasma samples from a retrospective cohort comprising 120 patients classified in 3 levels of severity according to the WHO ordinal scale. We found an association between the levels of pancreatic damage biomarker lipase and to a lesser extent the pancreatic amylase in COVID-19 patients according to severity of infection. Furthermore, these correlated with inflammatory biomarkers typically used in the clinic highlighting a relevant COVID-19-related pancreatic pathogenesis. Blood levels of pancreatic enzymes reflect a balance between the production and clearance of these enzymes. A robust or even a moderate rise in blood levels of lipase and amylase is usually associated with pancreatic injury^22,26^. Here, we report a 3- to 4-fold increase between critical vs severe subgroups in the proportion of patients with elevated plasmatic pancreatic lipase and amylase, respectively. The increased values are associated with histopathological changes and indicate the presence of an injury of the pancreas in COVID-19 patients that might be caused directly by the cytopathic effect triggered by the local SARS-CoV-2 replication. Of note, we cannot dismiss the possibility of an indirect cause for the pancreatic injury such as systemic responses to respiratory failure or the harmful immune response induced by SARS-CoV-2 infection, which also led to damages in multiple organs^35^. Interestingly, our MR study suggests a causal relationship between higher plasma levels of pancreatic amylase and severity of COVID-19. We identified significantly associated single nucleotide polymorphisms (SNPs) for each enzyme and performed a clustering analysis of these variants. A threshold of r2 < 0.001 was applied using the 1000 Genomes Project European reference panel, but a less restrictive threshold of 0.1 could be considered for future studies. Furthermore, further research is mandatory for the validation and confirmation of these findings in different populations and cohorts, as well as to perform the analysis in PLAC8, as a possible link between SARS-CoV-2 infection and up-regulation of PLAC8 expression in lung epithelial cells.

A previous CRISPR-Cas9 screening identified PLAC8 as an essential host factor required for SARS-CoV-2 lung infection^29^ and SADS-CoV swine infection^15^. Since a role of PLAC8 has been proposed in pancreatic tumorigenesis^11,12^, we investigated the contribution of this protein to the COVID-19-linked pancreatic pathophysiology. Analysis of pancreatic tissue from COVID-19 deceased patients showed substantially elevated PLAC8 levels as compared to uninfected controls (p<0.05). These findings are in line with robust PLAC8 expression documented in lung epithelial cells as demonstrated by single cell RNA-seq analysis^14,36,37^. Furthermore, PLAC8 is prominently expressed in the perturbed pancreas, especially in initial events of pancreatic cancer^10–12^.

To study the direct role of PLAC8 in SARS-CoV-2 infection, we generated PLAC8 knock-out by two different sgRNA CRISPR/Cas9 in a panel of human pancreatic cancer cell lines. We confirmed the effect of PLAC8 loss-of-function using full SARS-CoV-2 infectious virus inoculum from Wuhan-1 and BA.1 strains. Consistently, PLAC8 abrogation resulted in a substantial reduction in the number of infected cells in both KO cell lines tested. Importantly, evaluation of the replicative capacity reflected by infectious viral titers in the respective supernatants showed a decreased efficiency of viral production upon PLAC8 deletion. These results demonstrate for the first time the requirement of PLAC8 for SARS-CoV-2 viral replication in human tissue and are in accordance with previous observations addressing the swine acute diarrhea syndrome coronavirus (SADS-CoV)^15^ and SARS-CoV-2^14^. This further reinforces our findings and supports the potential of PLAC8 as a pan-CoV therapeutic target.

In conclusion, our data confirm the human pancreas as a SARS-CoV-2 target with plausible signs of injury unveiling the measurement of pancreatic enzymes with prognosis value and demonstrating that host factor PLAC8 is required for SARS-CoV-2 pancreatic infection defining new target opportunities for COVID-19-associated pancreatic pathogenesis.

## Author contributions

Conducted experiments: L I-G, SH, M L-D, D L-G, AM G-V, P G, S A-P, CC, JA M-G, CS

Investigation: CC, E C-B, APU, GB,

Designed experiments: SH, M L-D, T FE B, I F-C, CS, CB

Provided funding: CS, AK, CB

Writing, reviewing and editing: SH, M L-D, T FE B, I F-C, CS, CB

## Acknowledgements

We thank Prof Carlos López Otín for helpful comments. We thank Pau Pericàs-Pulido from PRISIB for technical assistance. We thank Dr Catalina Perello-Reus for helpful comments and assistance. We thank Victoria E Cano-Garcia from IDISBA Biobank for her assistance with sample processing. We thank Luis Enjuanes, at CNB-CSIC, for providing us with Vero E6 cells and with the Wuhan-1, and BA.1 strains of SARS-CoV-2. We thank Dr Marcos Malumbres (CNIO) for providing SUIT-2 cell line.

We thank Uta Lehnert for technical assistance. The imaging facility at the CMCB Technology Platform at TU Dresden is thanked for its assistance with imaging. The German Registry for COVID-19 Autopsies (DeRegCovid) is thanked for giving us access to the tissues from Regensburg.

This work was funded by the Instituto de Salud Carlos III (ISCIII) and Programa de Investigación en Salut—ISCIII (PI20/01267), by Fondo COVID19-ISCIII (COV20/00571), by Fundació La Marató (MARATÓ 167-C-2021 51), co-funded by the European Union and through the Programa Miguel Servet (MS19/00100) co-funded by the European Regional Development Fund/European Social Fund, “A way to make Europe”/“Investing in your future” (CB); TECH fellowship program, Impost turisme sostenible/Govern de les Illes Balears (TECH19/03) (LI-G);. Ministerio de Ciencia e Innovación MCIN/AEI /10.13039/501100011033 / FEDER, UE (PID-2021-123810OB-I00) and the European Commission – NextGenerationEU (Regulation EU 2020/2094), through Spanish National Research Council (CSIC)’s Global Health Platform (PTI Salud Global, CSIC-COV19-012/012202020E086) (MLD); “. The project that gave rise to these results received the support of a fellowship from “la Caixa” Foundation (ID 100010434). The fellowship code is LCF/BQ/DR22/11950020, awarded to DLG (DLG); Ministerio de Ciencia e Innovación (RYC2021-031776-I) (APU), the Deutsche Forschungsgemeinschaft (DFG, German Research foundation) project numbers 314061271 and 288034826 (CS). Ulm University Hertha Nathorff and Medical Scientist Program funding (SH).

## Data sharing

Any additional information required to reanalyze the data reported in this paper is available from the lead contact upon request.

