## Supplemental Figures for "Host factor PLAC8 is required for pancreas infection by SARS-CoV-2"

A

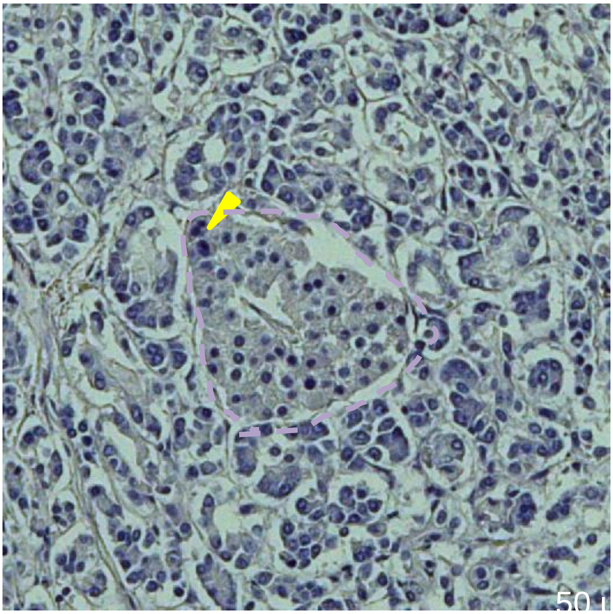

B

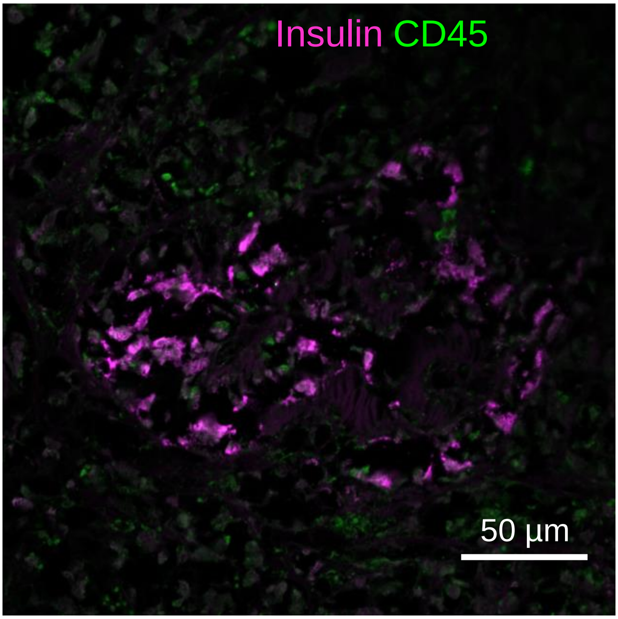


**Fig S1. Islet damage in the pancreas of COVID-19 patients.** A) H&E staining showing shrinkage of an islet with polyploid cells indicated with a yellow arrowhead. B) Immunofluorescence staining shows an islet with a low number of insulin-producing β-cells. Furthermore, infiltration with CD45 positive immune cells can be observed in both the exocrine and endocrine pancreas

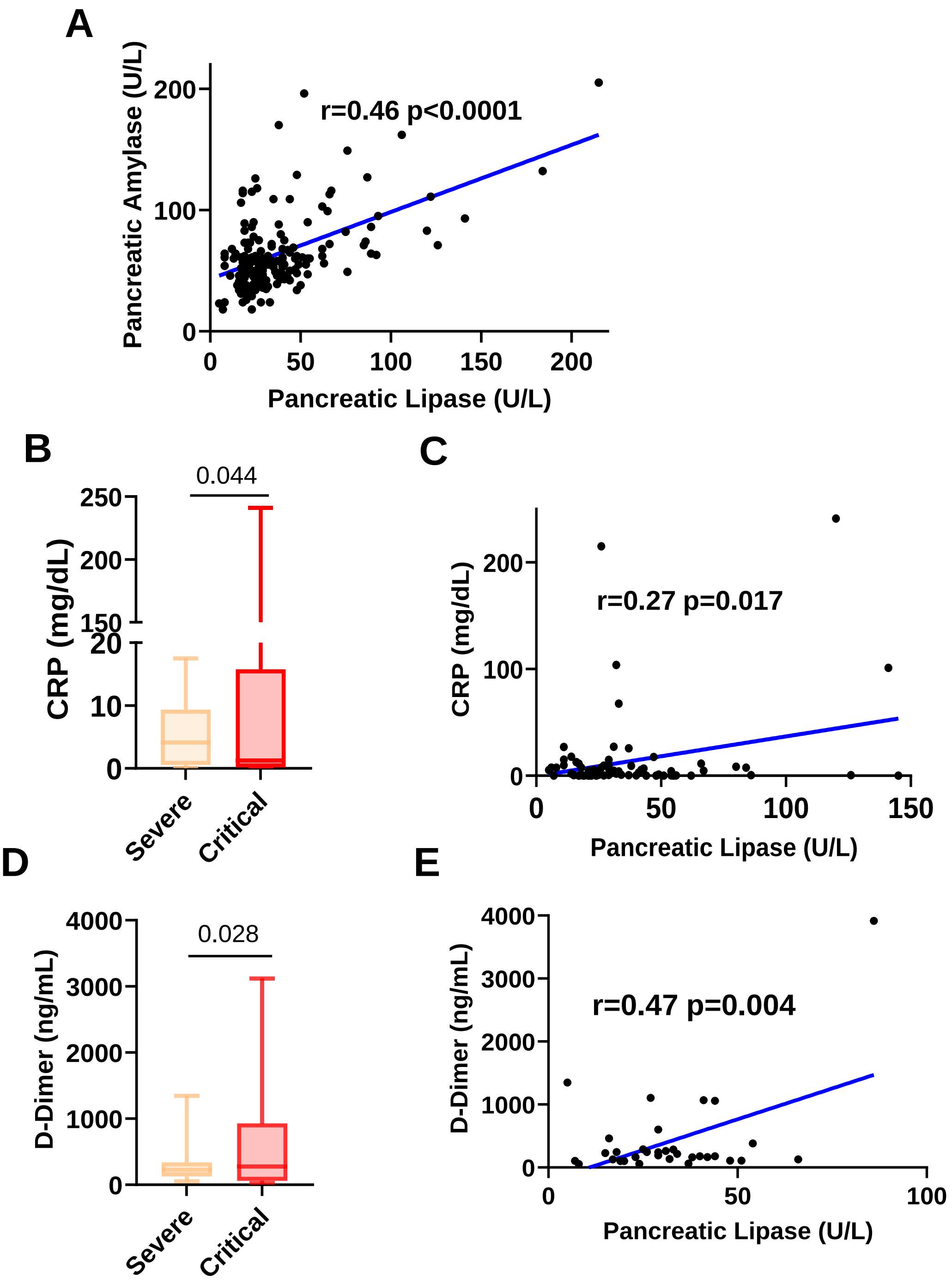


**Fig S2.- Anaysis of pancreatic enzymes and inflammatory plasmatic markers**

A) Correlation between plasmatic levels of pancreatic Amylase and Pancreatic Lipase (PNLIP) B) Plasma levels of COVID-19 severity biomarker C-reactive protein (CRP) across hospitalized COVID-19 patients C) Correlation between plasmatic levels of Pancreatic Lipase (PNLIP) and C-reactive protein (CRP) D) Plasma levels of COVID-19 severity biomarker D-Dimer across hospitalized COVID-19 patients E) Correlation between plasmatic levels of Pancreatic Lipase (PNLIP) and D-Dimer

**A**

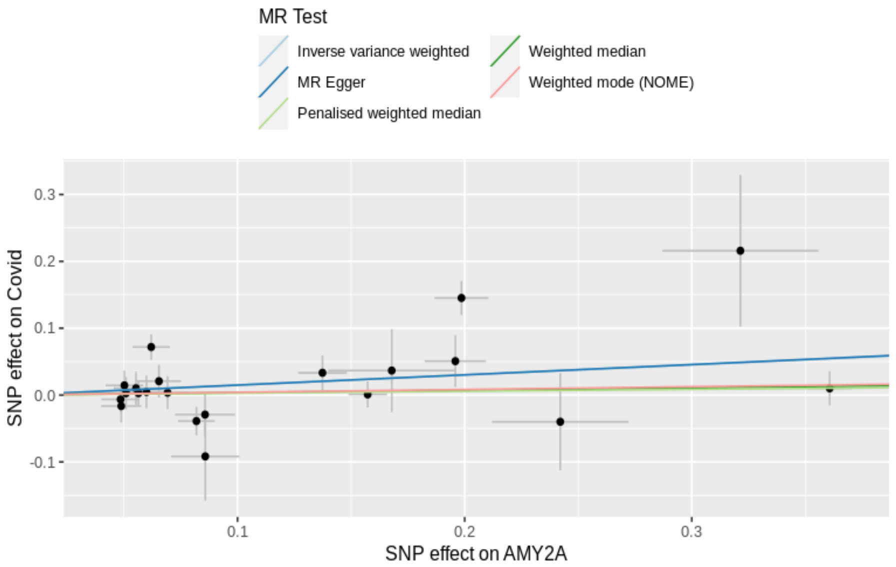


**B**

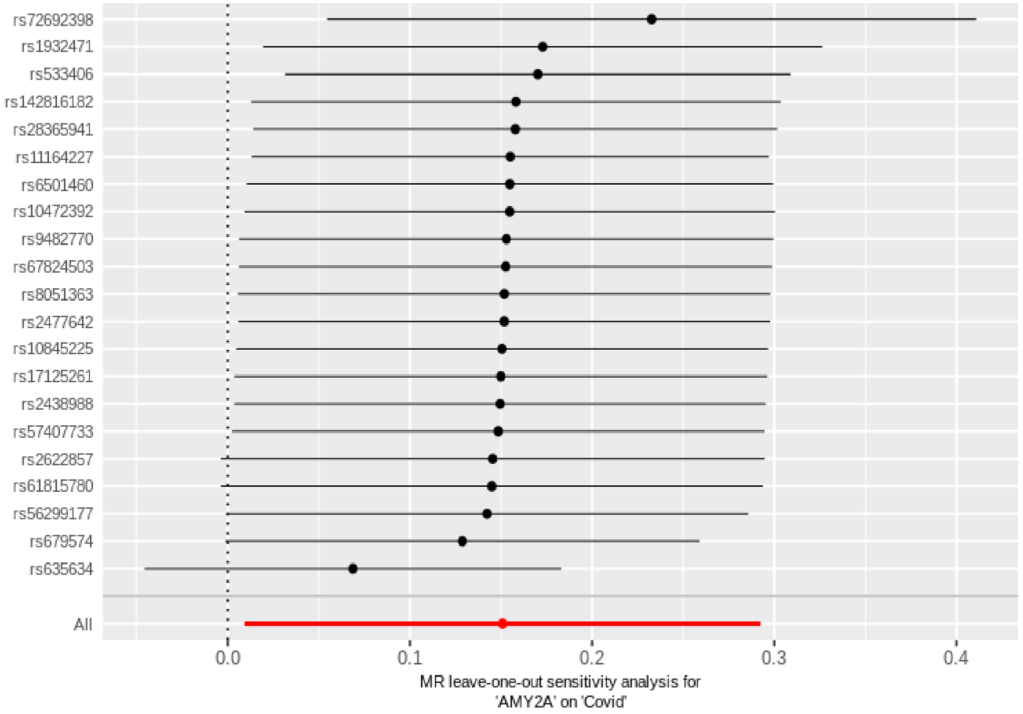


**Fig S3.- Mendelian randomization analysis of Pancreatic alpha-amylase (AMY2A) and COVID-19 severity**

A) Scatterplots of significant Mendelian randomization results. Analysis of AMY2A (Pancreatic alpha-amylase) and COVID-19 severity; SNP indicate single nucleotide polymorphism. B) Leave-one-out analysis for AMY2A on COVID-19 severity

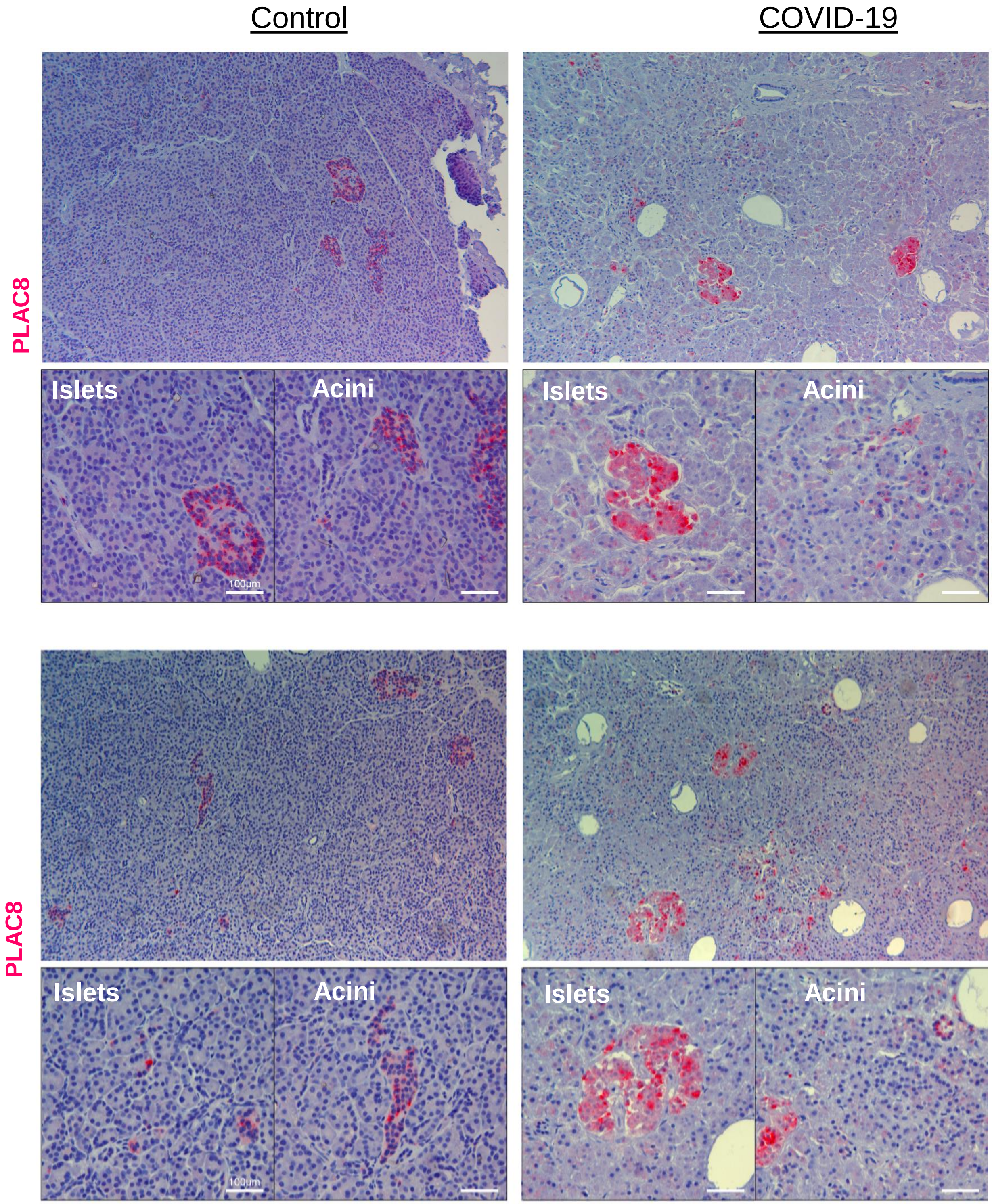


**Fig S4.- Representative images of PLAC8 staining in pancreas of non-infected control vs COVID-19 deceased donor**

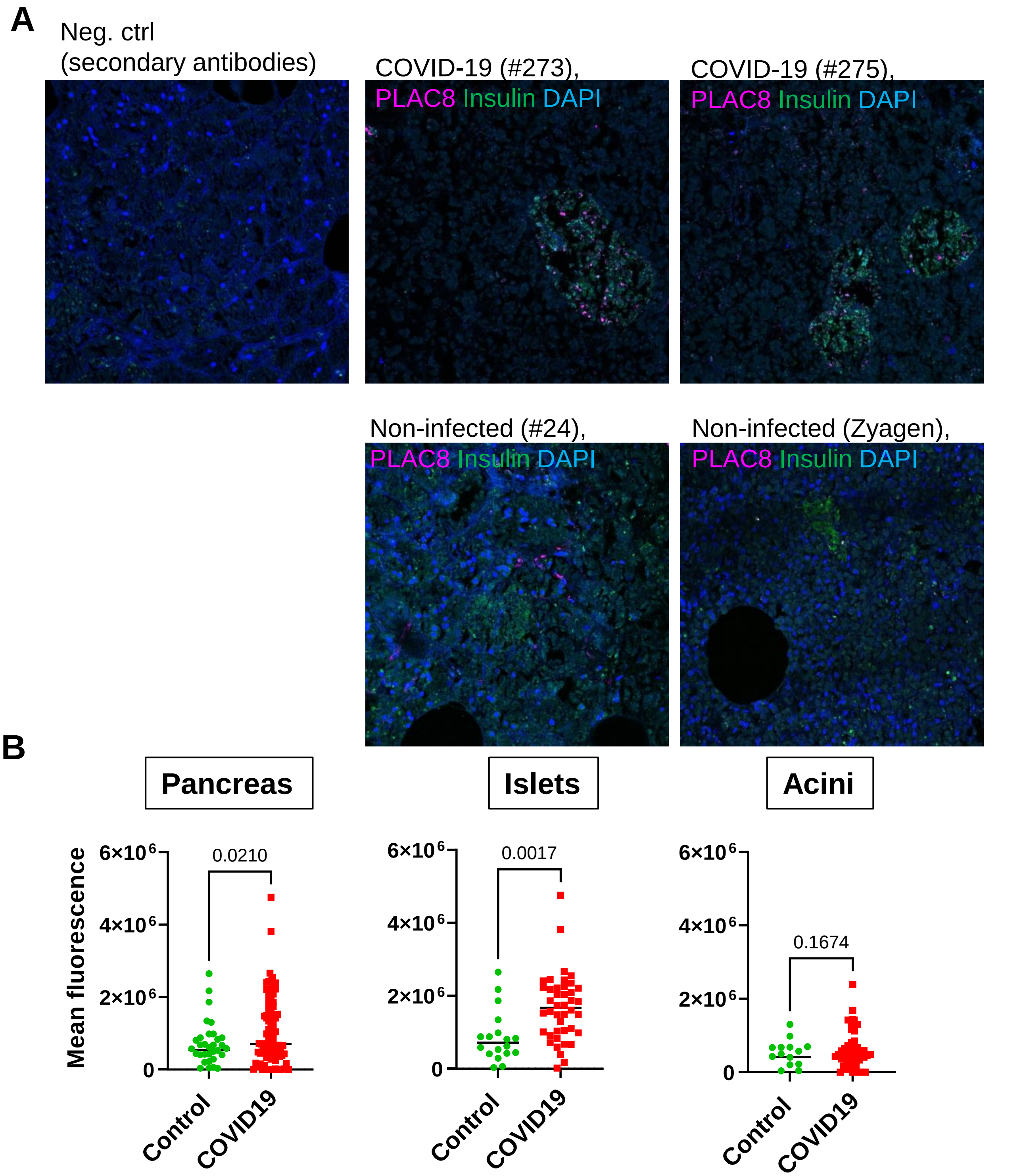


**Fig S5.- Immunofluorescence quantifications of PLAC8 positive cells using an independent series of COVID-19 patients (n=3) and non-infected donors (n=2).**

**A)**Representative images used in the quantification. A negative control (Neg ctrl) incubated with secondary antibodies was used to demonstrate the specificity of the staining**.**

**B)** Quantifications of mean fluorescence for whole pancreatic tissue (pancreas) and islets and acini compartments
